# Dynamic Adaptation of Default Mode Network in Resting state and Autobiographical Episodic Memory Retrieval State

**DOI:** 10.1101/2022.08.30.505423

**Authors:** Garam Jeong, Marco Paolini

## Abstract

The default mode network is a brain network processing mental states featuring an internal representation of subjective experiences like autobiographical episodic memory retrieval and a resting state. If the default mode network is the common spatial domain processing such mental states, then the temporal domain might present the differences in the mental states. To detect adaptations in dynamics of a single brain network dependent on the mental states it processes, we suggested a novel analysis method called one-to-many dynamic functional connectivity analysis for fMRI. The analysis method assesses the variance in the partial correlations of a center that are time-windowed functional correlations of a brain region (a center) to the rest of the regions in a brain network, then compares the similarities in the directions of their major variance from the same or distinct mental states. We applied one-to-many dynamic functional connectivity analysis to the default mode network and measured the similarity between the major variances of the partial correlations from three autobiographical episodic memory retrieval states and a resting state. If the major direction of the variance is a configuration presenting the mental states of the brain network, we expect to see the high similarity for the same mental states and less similarity for the distinct mental states. To test our hypothesis with the new analysis method, we chose a single subject who is a late blind with advanced mental imagery ability. The results showed that the dynamic adaption in the default mode network in the two mental states could be well depicted when the posterior cingulate cortex is the center in this single case. Furthermore, we could observe that the weight of the correlation between the posterior cingulate cortex and the right parahippocampal cortex varied mostly and therefore its contribution to the dynamic adaptation was relatively higher than the other correlations.

## 1 Introduction

For the last few decades, research on human brains has found that some brain regions are highly active when they are at rest without performing external tasks (Fox et al. 2005; Greicius et al. 2003). These brain regions include the posterior medial and the inferior lateral part of the parietal lobe and the medial prefrontal cortex in general. Resting-state functional connectivity analysis found that their activities are distinctly correlated (Biswal et al. 1995; Fox et al. 2005; Raichle et al. 2001). Because of these reasons, the set of brain regions was named the default mode network (DMN). The functions of the DMN are an ongoing research topic. Meta-analysis revealed that the DMN is highly active not only in a resting state but also in task-related mental states like episodic memory retrieval tasks, episodic future thinking tasks, theory of mind tasks, or navigation tasks (Spreng et al. 2009; Spreng & Grady 2010). All these task-related mental states construct internal mental imagery and focus on the imagery.

One of the suggested functions of the DMN regarding the commonalities between different brain states is “self-projection” (Andrews-Hanna et al. 2014; Buckner & Carroll 2007; Davey et al. 2016). Self-projection explains that the DMN processes the frame of reference of cognition formed with subjective experiences. Hence, these cognitive activities happen internally regardless of sensory inputs from outside. For example, one can listen to a bird song without subjective experiences by self-projection. However, retrieving memory related to the bird song requires self-projection. Another interesting suggestion for the possible functions of the DMN is “scene construction” (Hassabis & Maguire 2007). The previously mentioned mental states in meta-analysis commonly require mental imagery of scenes. Sensory imageries from memory are ingredients to form and maintain such scenes. The DMN might be the spatial domain to process such mental imagery. The self-projection and scene construction stress two pivotal aspects of the mental states mentioned by meta-analysis. One is the differentiation from perceptions generated with spontaneous sensory inputs, and the other is the creation of a coherent form of mental imagery. We summarize the two aspects as internal representations of subjective experiences in this paper.

The autobiographical episodic memory retrieval (AEMR) state is a mental state realizing these two suggested functions of the DMN (Conway 1996; Tulving 2002). Autobiographical episodic memory has an order between episodes (scenes) abstracted from subjective experiences in the past (Tulving 1972; Wheeler et al. 1997). The AEMR state asks subjects to retrieve successive scenes of their experiences in the past. Both self-projection and scene construction are keys for retrieving autobiographical episodic memory. In contrast to the AEMR state, a resting state is a mental state allowing the subject to drift in subjective perspective without the order of mental imagery or the need of picturing specific contents from past experiences. As a result, a resting state means often spontaneous mind wandering (Hurlburt et al. 2015). When the DMN is the spatial domain of neuronal substrates hosting the two mental states, could it be possible to distinguish dynamics of the DMN in the two mental states? To explore this idea, we compared the dynamics of the DMN in the two mental states.

### Dynamic Functional Connectivity

Functional connectivity analysis with fMRI measures correlations between the activities of two brain regions for the total measurement duration of BOLD signals (Friston et al. 1993). Research on functional connectivity has defined brain networks with and without tasks based on the correlations between brain regions across the whole brain (Allen et al. 2014; Shirer et al. 2012; van den Heuvel & Sporns 2013). Nevertheless, correlations in and between brain networks might change over time as cognitive activities do. Dynamic functional connectivity (DFC) is an analysis method to measure the changes in correlations over time (Allen et al. 2014; Chang & Glover 2010; Hutchison et al. 2013). Consequently, DFC research often focuses on the links between spontaneous alternations in cognitive activities and the corresponding adaptations in functional connectivity of brain networks (Bassett et al. 2011; Kucyi & Davis 2014; Saggar et al. 2018).

To analyze dynamics of a brain network and understand its implication, one should solve technical problems. There are two main concerns about analyzing BOLD signals from the perspective of network theory. One of the concerns is the high dimension of data. To understand dynamic adaptations of brain networks across the whole brain we need to consider a high number of variables and their dependency on each other. Thus, it is hard to catch the causality of such multivariate time series signals in connection to spontaneous cognitive activities. Further, when the connection between the observational method (fMRI BOLD) and the system under observation (neuronal substrates) is complex, we should interpret observations in our hands carefully. Previous studies on this issue have shown that the BOLD signals represent the neuronal activities of the brain reliably (Deco et al. 2011; Logothetis 2008). However, they contain also nonnegligible noises, and such noises could contaminate connectivity measurement (Keilholz et al. 2017; Tong et al. 2019). It could highly affect the results of DFC analysis when changes in correlation coefficients for the given time points are compared directly.

The concern over the complexity of the data due to its high dimension could be resolved by selecting well-defined features of brain networks. DFC studies in general analyze values of Pearson correlation coefficients of every pair in brain networks simultaneously (all-to-all). For n brain regions, therefore one considers *n*(*n*−1)*/*2 correlations for the given time points. To find corresponding regularities of the DFC of brain networks to mental states, researchers have hired various multivariate time series analyses using pattern recognition algorithms mainly (Calhoun et al. 2014; Hutchison et al. 2013; Leonardi et al. 2013; Preti et al. 2017). A big advantage of this approach is that one can grasp the dynamic organization of the functional networks across the whole brain regions. However, it is not easy to pick out a clear link between the dynamic organization and mental states of brain networks with many variables and potential noises summed up across the whole brain. For the cases like the DMN which is a well-defined spatial domain of mental states sharing a common functional feature, studying DFC of the single brain network only might be enough good to capture the link between dynamic organization in the brain network and the mental states.

We suggest a noble analysis method, one-to-many dynamic functional connectivity analysis (one-to-many DFC analysis) which is an analysis method for a single brain network. The steps of one-to-many DFC analysis are as follows: we choose a brain region, a center, from a single brain network then compute windowed Pearson correlation coefficients between the center and the rest of the brain regions in the brain network, which we call the partial correlations of the center. After that, the principal component analysis (PCA) is applied to the partial correlations of the center (Wold et al. 1987). Principal component analysis defines the principal components of the variance which is an orthonormal basis of the variance defined with the weight of each correlation of the center. The first few principal components are the direction explaining the most variance of the partial correlations. We hypothesized that the weight of each correlation of the center might be preserved for the same mental states and be varied between distinct mental states. That is, the direction of the major variance in the partial correlations described with the first or the second principal component is a configuration of the brain network corresponding to the mental state it processes. We can compare the principal components of the partial correlations of a center in the same and distinct mental states by measuring angles between them (Krzanowski 1979; Yang & Shahabi 2004). Consequently, if the weights of each correlation to the center represent a dynamic configuration of the single network corresponding to mental states of the brain network, we observe the differences or similarities in the major principal components for the distinct or the same mental states respectively.

The partial correlation of a center has *n* − 1 variables for a brain network with *n* brain regions. It includes not only a smaller number of variables but also conveys distinct aspects of a brain network compared to all-to-all DFC analysis. First, as already mentioned, the correlation coefficients measure the linear similarity between BOLD signals for the given time points, so it is susceptible to noises when we compare them directly. However, the directions of the variance in correlations are temporally abstracted properties of the correlations for the given time points. Therefore, spontaneous interruptions of noises could be evened out. Second, the principal components of the all-to-all correlations within a brain network are not equal to the sum of the principal components from the partial correlations of each center. Since the principal components are defined only regarding to the partial correlations of a center, it is possible to pick out which correlations of the center stay stable or change in various mental states.

In this study, we examined our hypothesis by applying one-to-many DFC analysis to the DMN of a single subject, since the analysis method is designed to study qualitative changes within an individual’s single brain network.

### Mental Imagery of the Blind

Because of the advanced visual system of humans, it is reckoned that visual stimuli dominate sensory inputs and are closely related to mental imagery (Hutmacher 2019; Pearson et al. 2015). Consequently, researchers have widely investigated the relationship between mental imagery and visual impairment to learn the mechanism of mental imagery (Cattaneo et al. 2008). The visual sensory input is spontaneous, and its sensory field is wide, so humans mainly rely on it to navigate spaces (Landau et al. 1981; Schinazi et al. 2016; Thinus-Blanc & Gaunet 1997). Some research on blind people shows that congenitally and late blind people can perform spatial navigation tasks as much as blindfolded sighted people can, although they are visually impaired (Schinazi et al. 2016; Tinti et al. 2006). At the same time, it found differences between the blind and the sighted people also. The size of sensory fields for auditory and tactile perception is smaller than those of the visual field. Hence, auditory and tactile perceptions contain local information and are naturally serial (Alain et al. 2001; Loomis et al. 1991; Röder et al. 2001). In contrast, the visual field offers global and local information simultaneously (Newell et al. 2005). Such a difference in sensory inputs leads blind people who prefer to employ auditory and tactile perceptions to process mental imagery through relative locations of objects in series (Iachini et al. 2014; Noordzij et al. 2006; Pasqualotto et al. 2013; Ungar et al. 1995). Accordingly, research on the serial memory of blind people shows that they could perform serial memory tasks better than sighted people do (Amedi et al. 2003; Postma et al. 2007; Raz et al. 2007; Röder et al. 2001). It might reflect their familiarity with the serial process. In addition to that, in the case of late blind people, activities of the visual cortex, including the primary visual cortex and the medial temporal lobe, were observed from mental imagery tasks (Amedi et al. 2003; Bridge et al. 2012; Goyal et al. 2006; Raz et al. 2005). It implies that late blind people can transfer the local and serial sensory inputs into global visual imagery. In short, the late blind people have advanced auditory and tactile perceptions contributing to local information and visual memory capturing a global structure of scenes by transferring local information into global visual imagery.

Concerning these facts, we assume that the late blind people have the advantage of having vivid mental imagery. In our study, we compared dynamics of neuronal activities at the network level with a single case of a late blind subject in two mental states, an autobiographical episodic memory retrieval state and a resting state. Following the ample studies on the late blind people summarized above, they have an advanced ability to retrieve serial memory fluently using vivid mental imagery that is closely related to episodic memory retrieval ability. Therefore, studying a late blind subject might facilitate catching the adaptation in dynamics of neuronal activities of the two mental states: the episodic memory retrieval state and the resting state. Furthermore, the late blind female subject aged 78 (called B.L.) who participated in this experiment previously attended an fMRI mental imagery study and the result showed that she could visualize vividly her episodic memory (von Trott zu Solz et al. 2017). We collected the fMRI BOLD signals of the late blind subject in the two mental states and analyzed dynamic functional connectivity in the DMN using one-to-many DFC analysis.

## 2 Material and Method

### 2.1 Experiment

B.L. had enucleation of both eyes in her adolescence due to infections. She had used the Braille system since her twenties. She had neither known brain damages nor neuronal diseases at the time she attended the experiment. In a previous experiment that she attended, her primary visual cortex activated highly when she was touching familiar geometric objects like a cylinder (von Trott zu Solz et al. 2017). It might imply that she had an advantage of retrieving episodic memory as summarized above. In addition to that, she had a morning routine that she could easily retrieve and state verbally with the consistency in the context. We asked the subject to retrieve her morning routine, which is a type of autobiographical episodic memory such that the subject could freely retrieve it without external cues (Conway 2009; Irish et al. 2011; Shirer et al. 2012). First, the subject described her morning routine verbally, then she repeated it three times in silence. After the behavioral experiment, the fMRI BOLD signals of the subject were collected while she was retrieving her morning routine and resting. The study was approved by the local ethics committee of the university’s medical faculty.

#### 2.1.1 Behavior

Before an fMRI session, the subject verbally described her morning routine and the description was recorded as an audio file. After the verbal statement, she retrieved the same morning routine three times in silence. The subject cued the beginning and the end of task performance verbally. An experimenter recorded the duration of each trial based on the cue.

#### 2.1.2 fMRI

We scanned the T1-weighted structural image with high resolution then the EPI sequences functional images. A foam cushion was used to minimize the effect of head movement. In the first fMRI run, B.L. was asked to rest without focusing on any specific thoughts or mental imagery. After the resting state run, she repeated the retrieval of her morning routine three times, and a short pause (less than a minute) was given in between fMRI trials. An experimenter cued the beginning and end of each run. The subject listened to the cue through a speaker in the MRI room. All four trials were 7 minutes 40 seconds following the result of the behavioral experiment. Imaging was performed with a 3T MRI clinical scanner (Phillips Healthcare, Ingenia). A T1-weighted 3D structural image was acquired in sagittal orientation. Its arrangements are as follows: repetition time (TR) = 8.2 ms, echo time (TE) = 3.76 ms, flip angle (FA) = 8°, number of slices = 220, matrix = 256 × 256, 1mm isotropic voxel size. Functional images were acquired after the T1-weighted anatomical reference scan using an echo-planar image (EPI) sequence. Its arrangements are as follows: repetition time (TR) = 2500 ms, echo time (TE) = 30 ms, flip angle (FA) = 90°, number of slices = 48, number of volumes = 180, matrix = 144 × 144, 3 mm isotropic voxel size, ascending sequential acquisition in axial orientation.

### 2.2 Analysis

#### 2.2.1 Data Preprocessing

Acquired T1-weighted anatomical images and EPI sequences were preprocessed by the default pre-processing pipeline of the CONN toolbox, which is a toolbox for SPM 12. The first six functional images were discarded, and 174 functional images (7 minutes 15 seconds) were used for further analysis. The preprocessing pipeline includes the realignment and unwrap, slice timing correction, outlier detection using ART, direct segmentation, normalization to MNI space, and functional smoothing. Realignment and unwrap were done to the first scan using b-spline interpolation. Slice timing correction was done to match the slice at each TR/2. Images showing their frame-wise displacement above 0.9mm due to head motion or the deviation of the global BOLD signal above five standard deviations were marked as outliers and scrubbed. Segmented functional and structural data were normalized to MNI space with 2 mm and 1 mm isotropic voxels respectively using 4th order spline interpolation. Functional smoothing was done with a Gaussian kernel of 6mm full width half maximum. To remove the slow physiological noises, the high pass filter with a 0.008 Hz threshold was applied.

#### 2.2.2 Regions of Interest

We employed group ICA of CONN toolbox and defined the DMN of the subject from images of the four fMRI trials. First, fast ICA with G1 contrast function abstracted the 40 spatial independent components after dimensionality reduction (dimension=64) (Calhoun et al. 2001; Hyvarinen 1999). We picked one of the components as the DMN by comparing spatial correlation and spatial overlap to the predefined DMN offered in CONN toolbox. The predefined DMN is defined with ICA of the resting-state data (n=497) of the Human Connectome Project (Lab 2017). The chosen independent spatial component contained the left and right angular gyrus (AG), the left and right parahippocampal cortex (PHC), the retrosplenial cortex (RSC), the precuneus (PRC), and a minor part of the posterior cingulate cortex (PCC) representing the posterior subsystem of the DMN. Additionally, we selected one more independent spatial component that could be regarded as the anterior subsystem or the midline core subsystem of the DMN mainly consisting of the PCC and the mPFC. Research on the DMN found that the DMN consists of a few subnetworks that may alter in number or appearance dependent on the study design and analysis methods (Andrews-Hanna et al. 2010; Damoiseaux et al. 2006; Davey et al. 2016; Mancuso et al. 2022). For that reason, we chose eight brain regions from two independent components in our study (Table 1). We extracted ROI marks of the PCC and the mPFC from an independent component with a threshold z-score ≥ 3, and the rest in another independent component with a threshold z-score ≥ 4 except for the L/R PHC using CONN toolbox. We chose a threshold of the z-score for the L PHC and R PHC as 2.5 to include a nontrivial number of voxels.

**Table 1:** MNI coordinates

|  | x,y,z | Number of Voxels |
| --- | --- | --- |
| L AG | -40, -75, 22 | 517 |
| R AG | 40, -71, 26 | 510 |
| L PHC | -29, -38, -17 | 86 |
| R PHC | 22, -38, -16 | 41 |
| PCC | 0, -53, 25 | 420 |
| mPFC | 0, 57, -7 | 847 |
| PRC | 0, -60, 42 | 539 |
| RSC | 2, -60, 16 | 765 |

Coordinates of the center for regions of interest in the MNI space.

#### 2.2.3 One-to-All Dynamic Functional Connectivity

One-to-many DFC analysis considers only correlations between a chosen center and its correlations to other brain regions in the same brain network at once. In this study, the DMN contains eight brain regions. We chose each of them as a center of one-to-many DFC at once. First, we calculated Pearson correlation coefficients of the partial correlations of a chosen center using sliding window method, then recentered them to have the zero mean to consider only the variance (Chang & Glover 2010). When we apply PCA to the time series of the seven partial correlations of the center in the DMN, it determines an orthogonal basis of the variance called the principal components. The first principal component is a unit vector indicating the direction where the most variance in the partial correlations of the center lies, the second most variance lies in the direction of the second principal component that is orthogonal to the first principal component, and so on. The biggest advantage of using PCA is the first few principal components explain the most variance of the multivariable system. If the first or the second principal component from the three trials of the AEMR state resemble each other, then one can say that the partial correlations of a chosen center have the high similarity in the direction encoding their major variance. On the contrary, one could expect that the first or the second principal components from the three AEMR trials and a resting state trial are dissimilar when it is indeed a configuration of the DMN corresponding to the mental state that it processes. We chose mainly the first principal components for most of our analysis. In the case of the second principal component of a trial explaining equivalently large variance in the partial correlations compared to the first principal component, we chose the one having a smaller angle to the chosen principal components of the other trials (Table 2).

**Table 2:** Chosen principal components (the amount of explained variance it explains)

| PC(%) | PCC | mPFC | L AG | R AG | L PHC | R PHC | PRC | RSC |
| --- | --- | --- | --- | --- | --- | --- | --- | --- |
| AEMR1 | 1(66) | 1(61) | 1(54) | 2(31) | 1(43) | 1(46) | 1(49) | 1(39) |
| AEMR2 | 1(65) | 1(60) | 1(50) | 1(55) | 2(32) | 1(59) | 2(24) | 2(21) |
| AEMR3 | 1(51) | 1(53) | 1(49) | 1(50) | 1(37) | 1(46) | 2(25) | 1(37) |
| Resting | 1(62) | 1(61) | 2(28) | 1(49) | 1(65) | 1(64) | 1(52) | 1(42) |

**Table 3:** Angles between chosen principal components of each fMRI trial

| (°) | PCC | mPFC | L AG | R AG | L PHC | R PHC | PRC | RSC |
| --- | --- | --- | --- | --- | --- | --- | --- | --- |
| AEMR1-AEMR2 | 8.98 | 12.88 | 49.37 | 33.09 | 25.00 | 20.64 | 35.84 | 48.50 |
| AEMR2-AEMR3 | 16.64 | 15.42 | 10.71 | 18.17 | 49.64 | 30.75 | 29.40 | 52.64 |
| AEMR3-AEMR1 | 13.54 | 13.44 | 44.43 | 48.14 | 34.96 | 18.05 | 31.09 | 10.06 |
| Resting-AEMR1 | 21.80 | 13.25 | 58.80 | 14.39 | 6.53 | 25.58 | 24.83 | 35.41 |
| Resting-AEMR2 | 22.17 | 19.28 | 13.22 | 45.06 | 22.07 | 44.62 | 34.42 | 52.41 |
| Resting-AEMR3 | 33.58 | 20.26 | 16.83 | 60.55 | 34.01 | 55.70 | 17.73 | 29.97 |

The similarity of the chosen principal components from the four fMRI trials (three AEMR trials and a resting state trial) can be depicted as angles between them (Gower 1967; Krzanowski 1979). Loadings of a principal component, *L*, indicate the weights of each correlation of the partial correlations to the principal component. Let us call the first principal component of a center *C* from a trial *A* is *PC*_*A*_(1). Then *PC*_*A*_(1) = (*L*_11_,…,*L*_17_) means that the *L*_1*i*_ the weight from the correlation between *C* and the brain region *i* to the *PC*_*A*_(1). When *PC*_*B*_(1) is a first principal component of a trial *B* to the center *C*, the angle *θ*_*AB*_ between the two principal components is calculated as follows:

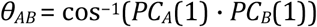

*PC*_*A*_(1) · *PC*_*B*_(1) is the inner product of *PC*_*A*_(1) and *PC*_*B*_(1) and the norm of each principal component is 1. The angle *θ*_*AB*_ tells how parallel the first principal component of the trial *A* and that of the trial *B* are. Similar methods to define the similarity of principal components from different samples were suggested previously (Krzanowski 1979; Yang & Shahabi 2004). When one of the first two principal components of the trial *A* is aligned parallel to that of the trial *B*, i.e., *θ*_*AB*_ is close to close to 0°, the chosen principal components are highly similar to each other. If the angle between chosen principal components is close to 90°, then the principal components are highly independent. Hence, we expect to see the small angles (close to 0°) between the chosen principal components of the partial correlations of the center *C* for three AEMR trials. Contrastingly, we expect big angles (close to 90°) between these principal components and the chosen principal component of the resting-state trial.

In this study, we employed the Pearson correlation coefficient to calculate the bidirectional and linear correlation between two brain regions and the sliding window method with the window width of 10 consecutive images (=25 sec with TR=2.5 sec). The weight of the window is constant so that the window has a rectangular shape. The window slid an image per step; therefore, 165-time points for 174 preprocessed images were used to calculate the variance in the partial correlations of a center. We applied PCA to the given distributions of partial correlations of a center for four trials each.

An example with a brain network of 4 ROIs. When *C* is the center, the correlations between *C* and the other three ROIs are computed using the sliding window method. The principal components (PC) of the partial correlations of *C* show its distribution on an orthonormal basis. The first and second principal component explain the most variance in the partial correlations of *C*. Therefore, we chose one of two principal components from each trial to compare their similarity. The angle between chosen principal components, for example, *θ*_*AB*_, measures the similarity of the first principal components from trial *A* and trial *B*.

## 3 Results

### 3.1 Behavior

B.L. reported that she could retrieve the same morning routine for the retrieval trials outside and inside the MRI machine. The subject described her morning routine as a sequence of episodes. The term ‘episode’ is used here to refer to events that happened in a specific location. Between episodes, navigational descriptions to move to the next location appeared. As an example, a scene in her bedroom is as follows:

> I imagined lying in bed having just woken up. I then put my feet to the left, as usual. Out of the bed, got the shoes, that were right there. Get up. Making my way around the bed, then there is the door. Through the door into the corridor. To the right and immediately to the left again, there is the toilet, something one needs in the morning. [translation from German]

In the verbal statement, retrieval in silence, and trials in the MRI machine, she reported that the last scene of her morning routine was when she is waiting for her husband sitting at the breakfast table in the living room. 7 minutes 40 seconds was the mean of measured time for the three retrieval trials in silence (SD *<* 10 sec).

### 3.2 fMRI

The first three principal components of four trials explained over 80% of the entire variance in the partial correlations for each chosen center in the DMN. Criteria for selecting a principal component from a trial were as follows: we computed angles between the first two principal components of the partial correlation of a center from the three AEMR trials one by one, i.e. *θ*_*AB*_(*i,j*) for *A,B* = 1,2,3 and *i,j* = 1,2 is the angle between the *i*th and *j*th principal components of a trial *A* and a trial *B* respectively. Then we chose a *PC*_*A*_(*i*) having the smallest angle to a *PC*_*B*_(*j*) for *A* ≠ *B*. When a *PC*_*A*_(*i*) has the smallest angle to the chosen principal component of the trial *B* and a *PC*_*A*_(*j*)(*i ≠ j*) has the smallest angle to the chosen principal components of another trial, for example a trail C, then we chose the principal components having the smaller mean angle between three of them. Then we chose a principal component of the resting state trial having the smallest angle to the chosen principal components of three AEMR trials. If the two principal components of the partial correlations of the center from the resting state trial have the smallest angle to the trial A and the trial B each, then we choose the one giving the smallest mean angle between three from the AEMR trials and one from the resting state. As shown in Table 2, the chosen principal component of each trial is mainly the first principal component. When the second principal component was chosen from a trial, the variance it explains was relatively similar to the first principal component of the trial.

The mean angle between the chosen principal components of the three AEMR trials and that of all four trials including the resting state trial were calculated for each center. It assesses the similarity between the chosen principal components of the partial correlations of a center (Table 4). Also, we gauged the changes in the mean angle in the case of including the principal component from the resting state trial with the ratio between the two mean angles. The implication of the ratio is as follows: when the mean angle between three principal components from the AEMR trials is equal to the mean angle between four principal components including the chosen principal component of the resting-state trial, the ratio is 1. That is the chosen principal components are consistent for the two mental states. The chosen principal components of the three AEMR trials have higher similarity between them compared to that of the resting state trial when the ratio is smaller than 1. The smaller the more differences between the chosen principal components of the AEMR trials and the resting state trail. By contrast, the chosen principal components of the three AEMR trials have higher similarity to that of the resting-state trial than they do between themselves when the ratio is bigger than 1 (Table 4).

**Table 4:**
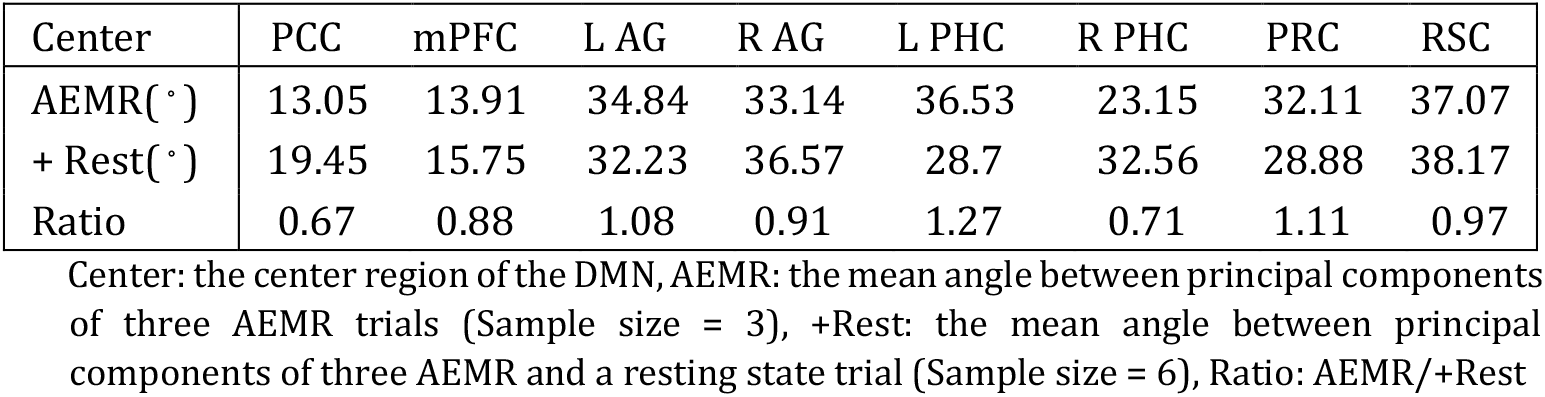
Mean angles between chosen principal components and the ratio between them

| Center | PCC | mPFC | L AG | R AG | L PHC | R PHC | PRC | RSC |
| --- | --- | --- | --- | --- | --- | --- | --- | --- |
| AEMR(°) | 13.05 | 13.91 | 34.84 | 33.14 | 36.53 | 23.15 | 32.11 | 37.07 |
| + Rest(°) | 19.45 | 15.75 | 32.23 | 36.57 | 28.7 | 32.56 | 28.88 | 38.17 |
| Ratio | 0.67 | 0.88 | 1.08 | 0.91 | 1.27 | 0.71 | 1.11 | 0.97 |

Center: the center region of the DMN, AEMR: the mean angle between principal components of three AEMR trials (Sample size = 3), +Rest: the mean angle between principal components of three AEMR and a resting state trial (Sample size = 6), Ratio: AEMR/+Rest

As shown in Table 4, the principal components from the partial correlations of the PCC center have the smallest mean angle (13.05°) between the three AEMR trials. In addition, its ratio to the mean angle of the four principal components of all trials (AEMR/+Rest = 0.67) was the smallest compared to the ratios of the other centers in the DMN. The small angles between the first principal components of three AMER trials indicate that the DMN relatively well preserved the weights of the partial correlations of the PCC to the chosen major principal components for the AMER trials. Furthermore, the ratio between the mean angle with the AEMR trials and the mean angle together with a resting state trial showed that the weights have relatively large dissimilarity in the two different mental states. In Figure 3, differences between the mean weights of the partial correlations of the PCC to the chosen principal components of the AEMR trials and that of the resting-state trial are presented. The largest change among the weighs of the partial correlations of the PCC was observed at the weight of the correlation between the PCC and the R PHC.

**Figure 1:**
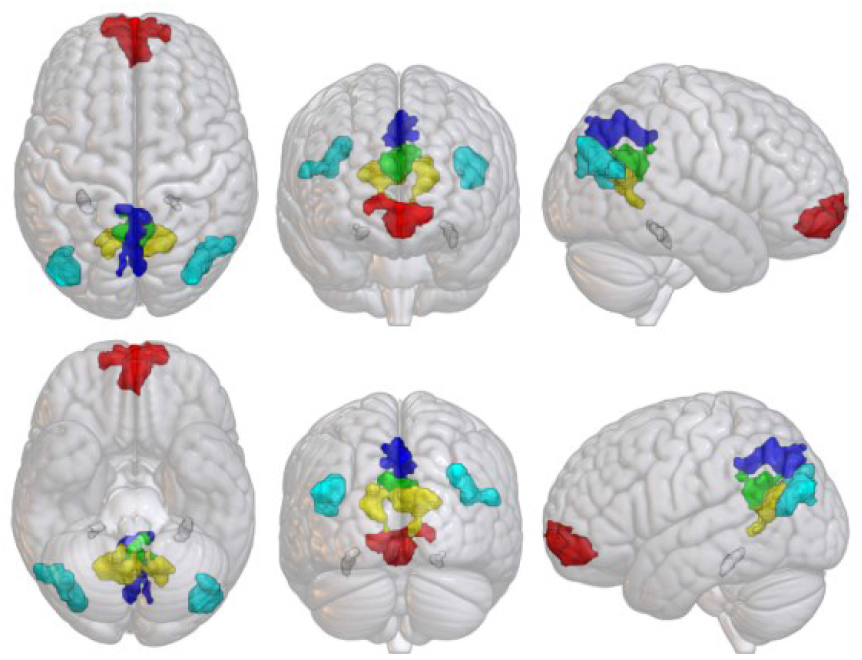
Regions of interest L/R AG – Cyan; L/R PHC – white; PCC - green; mPFC – red; PRC – blue; RSC-yellow The image was created by MRIcroGL.

**Figure 2:**
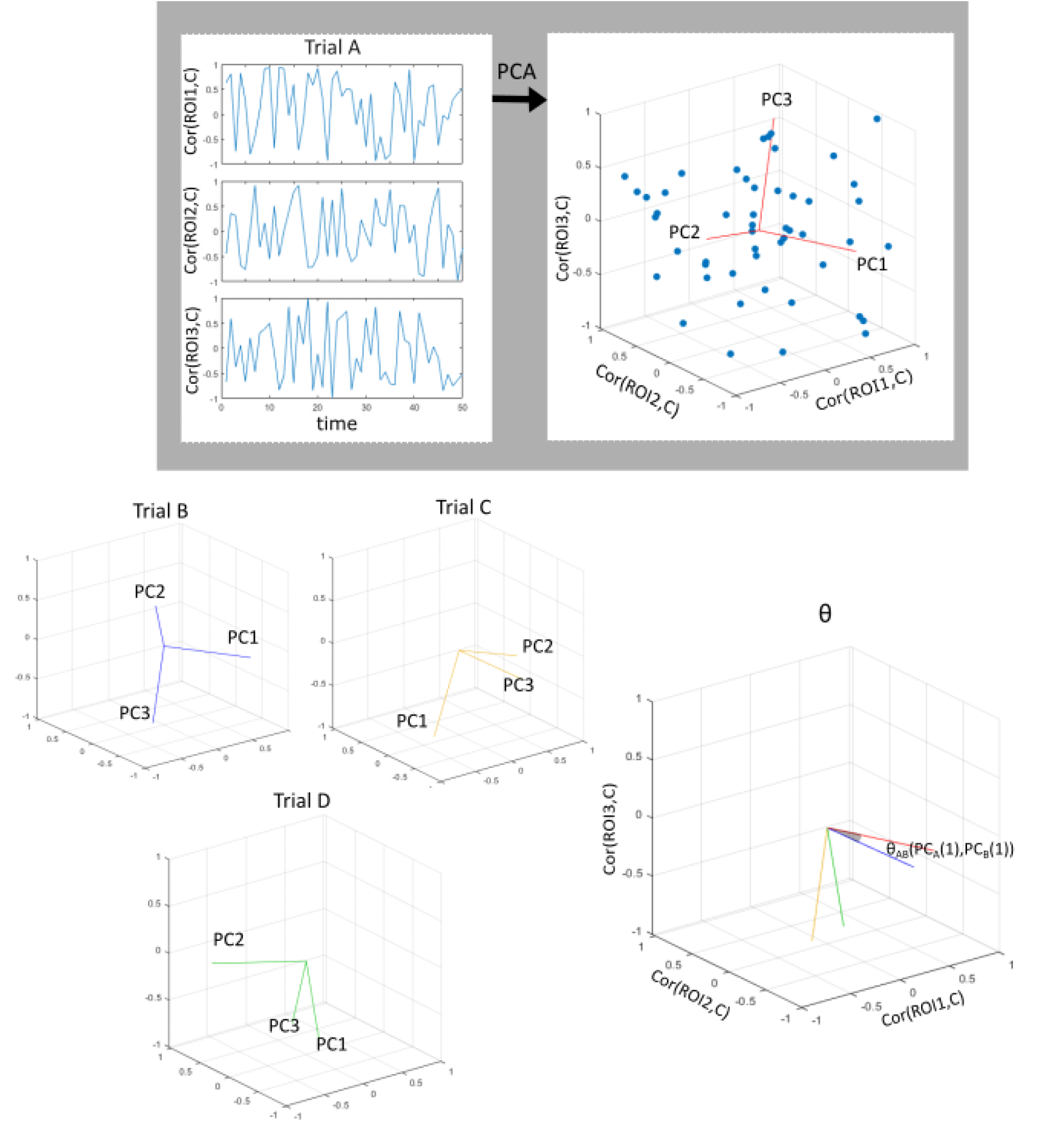
One-to-all dynamic functional connectivity analysis

**Figure 3:**
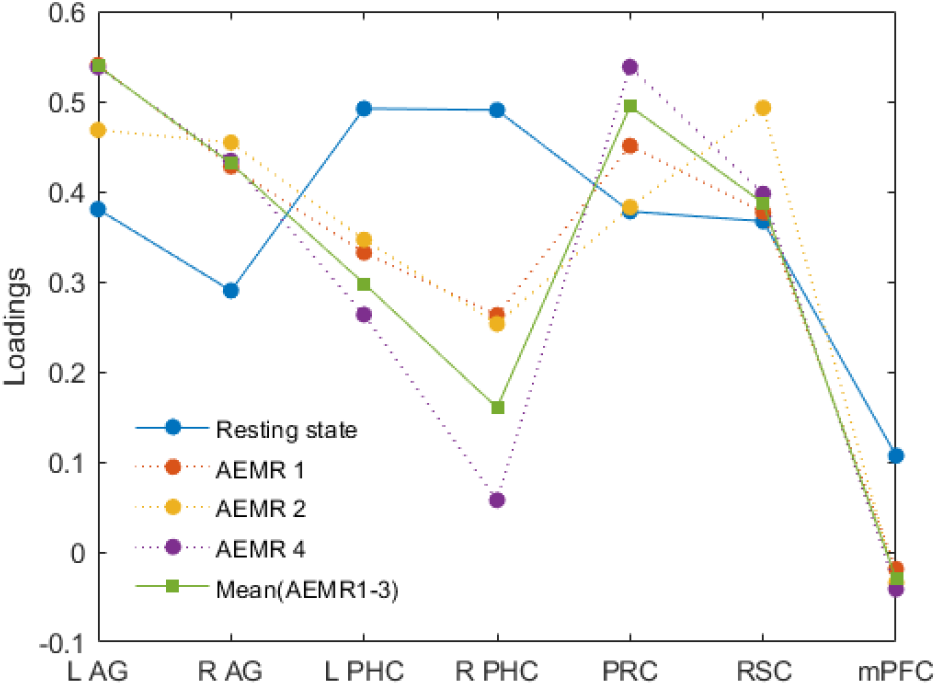
Loadings of the partial correlations of the PCC center

The chosen principal components of three AEMR trials with the mPFC center also had a relatively small mean angle (13.91°). However, in contrast to the case of the PCC center, its ratio to the mean angle together with the chosen principal component of the resting-state trial was relatively large (0.88). This result implies that the weights of the partial correlations to the chosen principal components were rather similar in both mental states compared to the PCC center case. For the case of the R PHC center, the ratio of the two mean angles is relatively small (0.71). Nevertheless, the mean angle between the chosen principal components of the AEMR trials (23.15°) is almost twice larger as that of the PCC center case. Except for the PCC, the mPFC, and the right PHC center, we could not find other brain regions either having a high similarity between the chosen principal components of the three AEMR trials or having clear dissimilarity between them and that of the resting-state trial.

## 4 Discussion

In this exploratory study, we aimed to investigate adaptations of the DMN in the two mental states, the AEMR state and the resting state, with a new analysis method called one-to-many DFC. For this purpose, we conducted the behavioral and fMRI experiment with a blind subject who has an advanced ability to envisage mental imagery. Given the verbal description of the morning routine from the behavioral experiment, we could see that the spatial navigation guided the streaming of the episodes mostly. Also, the subject could retrieve the morning routine in silence with a constant retrieval duration for each trial. These results from the behavioral experiments imply that the subject could retrieve the morning routine with regularity, which is a distinctive feature of the AEMR state compared to a resting state. In the fMRI experiment, dynamic functional connectivity of the DMN in the AEMR state and the resting state were analyzed by one-to-many DFC analysis. The new analysis method measured only the partial correlations of a center in the DMN at once then computed an orthonormal basis of the variance in the partial correlations using principal component analysis. The most parallelly aligned principal components among the first two principal components of each trial were chosen to compare their similarity by measuring the angle between them. In short, we suggested probing a dynamic configuration of the DMN, which is the known spatial domain of the two mental states by comparing the directions of the major variance in the partial correlations of a chosen center. The partial correlations offer specific information about a single brain network regarding a center which is the relation between a chosen center and the rest of the brain regions in the single brain network. Therefore, it is possible to see the adaptation of the single brain network in terms of changes in the weights of the correlations of the chosen center. In the case of the three AEMR trials, the results of one-to-many DFC analysis showed that the major variance of the partial correlations of the PCC center most similarly aligns with each other (13.05°) compared to the results of the other centers (Table 4). That is the configuration of the DMN defined as the weights of the partial correlations of the PCC to a chosen principal component was most consistent for the three AEMR trials. At the same time, we computed changes in the configuration of the DMN by measuring the ratio of the mean angle between the chosen principal components of three AEMR trials to the mean angle between all four chosen principal components including the one from the resting state trial. The result suggested that the principal components of the three AEMR trials align with that of the resting-state trial in the most distinct direction when the PCC is the center of the partial correlations (AEMR/+Rest = 0.67) (Table 4). Previous fMRI studies on the DMN showed that the PCC is the most reliable marker in the adaptation of the DMN by measuring changes in bold signals and correlations of the DMN (Davey et al. 2016; Xu et al. 2016). In our study, we could prove the consistent result with one-to-many DFC analysis by hiring a subject-driven experimental paradigm that is close to the natural cognitive activities (Shirer et al. 2012). Furthermore, we could pick out that the weight of the correlation between the PCC and the R PHC varied mostly so that it determines the dynamic configuration of the DMN corresponding to the mental state it processes.

Interestingly, the PCC is a well-known functional and structural hub of the brain networks (Achard et al. 2006; Buckner et al. 2009; Hagmann et al. 2008; Honey et al. 2007; Tomasi & Volkow 2011). To date, several studies on brain networks have speculated that hubs in human brains boost efficient communication between brain regions (Bassett et al. 2006; Bullmore & Sporns 2009; Rubinov & Sporns 2010; van den Heuvel & Sporns 2013). Following the idea of hubs of brain networks, it is reasonable to expect to see that connections between brain regions in a single brain network adapt themselves via its hubs. Correspondingly, the results of one-to-many DFC analysis on the DMN demonstrated that the DMN adjusts its configuration via the PCC. By contrast, the analysis results of the mPFC center, which is also one of the known functional and structural hubs of the brain, did not demonstrate the distinctive configurations in the two mental states (AEMR/+Rest = 0.88) compared to the PCC center case. We could therefore speculate that the local hub of the DMN mediating adjustment within the network is likely to be the PCC, but less likely to be the mPFC. Consistent with the implication, the anatomical and structural connections between the PCC and the other brain regions in the DMN are confirmed by cytoarchitecture and diffusion tensor imaging studies (Greicius et al. 2009; Vogt et al. 2006).

Besides the results of the PCC center case, the result of the analysis with the R PHC center demonstrated that the configurations of the DMN in the two mental states were relatively dissimilar, which was described by the small ratio between the two mean angles (AEMR/+Rest = 0.71). Unlike the PCC and the mPFC, the R PHC is not one of the known hubs in brain networks. In accordance with this, the chosen principal components of three AEMR trials of the R PHC center aligned less coherently (23.15°) compared to that of the PCC (13.05°) and the mPFC (13.91°). The results could be explained by the fact that the weight of the correlation between the PCC and the R PHC for the chosen principal components varied considerably in the two mental states. As we suggested in Figure 4, the loading of the R PHC to the PCC center varied around twice bigger as that of the other regions. A possible implication of this result is that the functional connectivity between the PCC and the R PHC takes a pivotal role in explaining the difference between the two mental states and it caused the relatively large changes in the mean angles for the R PHC center case too. One of the known functions of the PHC is related to the recognition of local scenes and contextual association (Aminoff et al. 2013; Epstein 2008; Maguire et al. 1998). Accordingly, the activity of the PHC might be the key to retrieving coherent mental imagery of scenes, in contrast to a resting state which does not require such a function. Moreover, studies on the lateralization of the medial temporal lobe (MTL) have suggested that the major function of the right MTL is related to spatial navigation (Golby et al. 2001; Miller et al. 2018; Spiers et al. 2001). In our study, the result from one-to-many DFC analysis together with the verbal statement of the subject, therefore, could be interpreted as follows: spatial navigation in the mental imagery of the subject might drive the substantial adaptation of the dynamic configuration of the DMN in the two mental states determined with the one-to-many DFC analysis.

**Figure 4:**
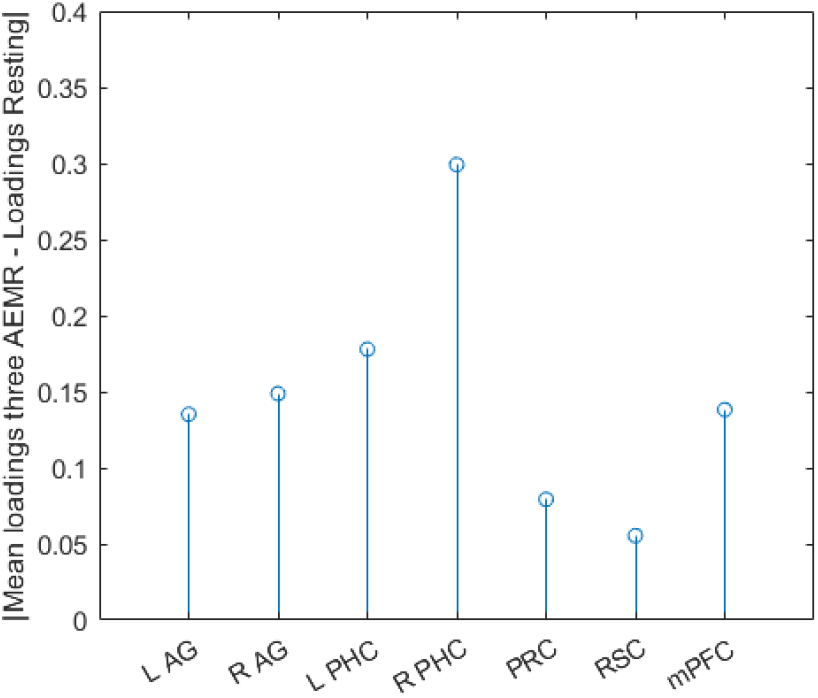
Differences in Loadings of the partial correlations of the PCC center

In this paper, we aimed to explore the new analysis method, one-to-many DFC analysis thoroughly. We took a single subject having an advanced ability to perform mental imagery tasks, thereby facilitating the validity of the analysis method. The results of the analysis suggested that the consistency in dynamics of the DMN in the same mental states and its adaptations in the distinct mental states could be described with one-to-many DFC analysis. Therefore, we would like to suggest that the weights of the correlations between the PCC and the rest of the brain regions in the DMN to the major direction of their variance form a reliable configuration to describe its dynamic feature corresponding to the mental states. Furthermore, the result could depict where the largest change in the configuration comes from.

However, being limited to a single subject, this study lacks generality. To achieve that, we conducted the same experiment with several subjects, which will be reported separately. We would like to close this section by pointing out an interesting possible extension of the present study. The configuration of a local brain network defined by one-to-many DFC analysis focuses only on linear properties of DFC of partial correlations. The nonlinear properties of partial correlations may reveal further instructive information about a single brain network.

## 5 Conclusion

It is believed that dynamic adaptation in brain networks reflects the streaming of mental states, which is the nature of humans. One-to-many DFC analysis may offer us a tool to describe an interesting dynamic feature of a single brain network that is preserved in specific mental states. This research has thrown up many questions in need of further investigation, like its application to other known brain networks in various mental states including psychological disorders and neurodegenerative diseases.

## Acknowledgements

We would like to thank Isolde Zondler and Dr. Urs Zondler for their financial support for this work.

